# Earthworms and plants can decrease soil greenhouse gases emissions by modulating soil moisture fluctuations and soil macroporosity in a mesocosm experiment

**DOI:** 10.1101/2023.07.28.550960

**Authors:** Pierre Ganault, Johanne Nahmani, Yvan Capowiez, Nathalie Fromin, Ammar Shihan, Isabelle Bertrand, Bruno Buatois, Alexandru Milcu

## Abstract

Earthworms can stimulate microbial activity and hence, greenhouse gas (GHG) emissions from soils. However, the extent of this effect in the presence of plants and soil moisture fluctuations, which are influenced by earthworm burrowing activity, remains uncertain. Here we report the effect of earthworms (without, anecic, endogeic, both) and plants (with, without) on GHG (CO_2_, N_2_O) emissions in a 3 month-greenhouse mesocosm experiment simulating a simplified agricultural context. The mesocosms allowed for water drainage at the bottom to account for the earthworm engineering effect on water flow during two drying-wetting cycles. N_2_O cumulative emissions were 34.6 and 44.8% lower when both earthworm species and only endogeic species were present, respectively, and 19.8% lower in presence of plants. The presence of the endogeic species alone or in combination with the anecic species slightly reduced CO_2_ emissions by 5.9% and 11.4% respectively, and plants presence increased emissions by 6%. Earthworms, plants and soil water content interactively affected weekly N_2_O emissions, an effect controlled by increased soil dryness due to drainage via earthworm burrows and mesocosm evapotranspiration. Soil macroporosity (measured by X-ray tomography) was affected by earthworm species-specific burrowing activity. Both GHG emissions decreased with top soil macropore volume, presumably due to reduced moisture and microbial activity. N_2_O emissions decreased with macropore volume in the deepest layer, likely due to fewer anaerobic microsites. Our results indicate that, under experimental conditions allowing for plant and earthworm engineering effects on soil moisture, earthworms do not increase GHG emissions and that endogeic earthworms may even reduce N_2_O emissions.

## Introduction

Soil invertebrates, by their feeding activity, movement through soil layers, and interactions with other organisms, strongly modulate organic matter dynamics, carbon and nutrient cycling, as well as water and gas fluxes [1–3]. Particularly important, earthworms promote soil fertility and plant productivity [4,5], with 25% increase in crop yield [6] in their presence. Conversely, the presence of earthworms increases CO_2_ emissions by 33% and N_2_O emissions by 42% [7], which is notable since N_2_O has 265 times the global warming potential of CO_2_ [8]. However, the microbial processes causing greenhouse gas (GHG) emissions are extremely complex, and involve numerous interactions between earthworms, microbial communities, plants, and soil water and aeration status [9–11]. Simultaneously exploring the complexity of these interactions and mechanisms mentioned above in experimental settings poses a challenge. Most existing studies have focused on scenarios without of plants and at constant soil moisture levels that favor earthworm and microbial activity [7]. This limits our understanding of how soil moisture fluctuations, modulated by earthworms and plants and their interactions, affect N_2_O and CO_2_ emissions [7,12].

Soil water content is a long recognized key factor, explaining up to 95% GHG emissions [13], as it drives microbial processes such as respiration, denitrification, and nitrification that produce GHG [9,14,15]. Indeed, soil water content determines gas and nutrient diffusion and hence the availability of oxygen, nitrate, ammonium and carbon to microorganisms, thereby modulating their activity. Under anoxic conditions, at high soil water content, N_2_O emissions are the highest, mostly produced by denitrification, while aerobic conditions favor N_2_O emissions by nitrification [16,17]. Similarly, a substantial body of evidence showed that carbon substrate limitation occurs in drier conditions, while oxygen limitation occurs under conditions close to water saturation, with optimal conditions for respiration, and hence CO_2_ emissions, at intermediate water contents [18,19]. Soil moisture fluctuations (e.g. drying-rewetting cycles) can also affect the proportion of nitrogen denitrified into N_2_O or N_2_, thus modulating the N_2_O/N_2_ ratio that will be emitted into the atmosphere [20,21]. Keeping constant soil moisture like in most existing experiments therefore limits these mechanisms from occurring.

As primary producers, plants control organic matter quantity and quality in soils [22]. Inputs of root-derived C substrates leads to high transient O_2_-demand and can cause suboxic microsites in the rhizosphere, thus favoring denitrification [9,23]. Conversely, plants compete with microbes for nitrogen acquisition, reduce soil water content by transpiration, modify soil porosity by root growth [24,25], and thus can change the preponderance of the controlling N_2_O emission processes (nitrification, denitrification) [9,26]. Simultaneously, earthworm ingestion of organic matter and mixing with the mineral soil and their mucus modify the accessibility and availability of nutrients and water to microbes, favoring denitrification and mineralization on the short term, and carbon stabilization on the long term [27–30]. On the other hand, earthworm burrowing through the soil profile increases soil aeration, water drainage and possibly soil drying [31–34], thus potentially creating conditions less favorable for denitrification and N_2_O emission [12,35]. The above mechanisms interact all together since earthworms promote plant growth via increasing nutrient availability to plants and by consuming roots [4–6]. To the best of our knowledge, no study investigated earthworm-mediated soil moisture variations effect of GHG emissions. Furthermore, the vast majority of existing experimental studies used micro/mesocosms with sealed bottoms, which impede water drainage and the modulation of water infiltration by earthworm burrows and root growth that would typically occur in realistic field conditions. Two studies looked at the impacts of soil moisture fluctuations on GHG in the presence of earthworms, although without plants, showed that cumulative N_2_O and CO_2_ emissions were reduced in the presence of earthworms [12,35].

Allowing earthworm burrowing activity to influence soil water content could help understand the varying effects of different earthworm species and ecological categories on earthworm-mediated GHG emissions. Indeed, lumbricid earthworms are broadly classified in three main ecological categories based on their feeding and burrowing characteristics: (1) anecic species that feed on surface litter by pulling it into permanent vertical burrows and create surface casts, (2) epigeic species that live, feed in surface litter, making very few non-permanent burrows and (3) endogeic species that live in the soil, feeding on roots and soil organic matter, and making numerous non-permanent burrows [36,37]. Hence, anecic earthworms are likely to have a stronger impact on organic matter redistribution and water fluxes. On the other hand, endogeic and epigeic earthworms may primarily influence organic matter redistribution in the soil and at the surface, respectively, with epigeic earthworms having a lesser impact on water fluxes. There is evidence that CO_2_ and N_2_O emissions depend on earthworm ecological category, with significantly higher emissions for the anecic group [7], but the net balance between mineralization and stabilization over time was reported to be highly variable within each category [38]. Additionally, whether this finding holds in the presence of plants and soil moisture fluctuations remains to be tested.

In this study, we assessed the impact of an earthworm treatment with four levels (one endogeic species, one anecic species, both anecic and endogeic species, and a control without earthworms), and a plant treatment with two levels (with or without a model grass species). We used a full factorial design (4 × 2 = 8 treatment combinates, with 7 replicates each), during a three months greenhouse mesocosm experiment simulating a simplified agricultural setup. The experiment involved simulating two drying- wetting cycles, and to facilitate the earthworm engineering effect on soil water infiltration and status, the mesocosms were designed to enable effective water drainage and escape via percolation. We measured weekly CO_2_ and N_2_O fluxes, aboveground plant biomass, litter cover and multiple soil parameters representing potentially relevant predictors of GHG emissions including soil nitrogen and water status, microbial biomass and respiration, denitrification potential and multiple metrics of soil macroporosity using X-ray tomography. We hypothesized that: 1) CO_2_ and N_2_O emissions will be lower in the presence of earthworms relative to controls as increased carbon and nitrogen mineralization will be offset by the drier and more aerated conditions due to earthworm soil engineering effect (burrowing) on water drainage, 2) plant presence will reduce N_2_O emissions due to nitrogen and water uptake but will increase CO_2_ emissions due to increased carbon substrates entering the soil via rhizodeposition, and 3) differences in N_2_O and CO_2_ fluxes among the two earthworm ecological categories are mediated by the burrowing patterns affecting soil (macro) porosity and water status, expecting lower emissions with higher macroporosity, as this increases the volume of aerobic and dryer sites.

## Materials and methods

### Soil and biological material

The soil, classified as a gleyic luvisol, was excavated from a field margin adjacent to a wheat- corn-alfalfa rotation at the EFELE experimental site (North West of France, 8°05′35.9”N, 1°48′53.1”W) belonging to the long-term observatories SOERE-PRO-network. Only soil from the upper 0-30 cm layer was used in this experiment. The soil is composed of 14.6% clay, 72.1% silt and 13.3% sand, with a pH of 6.14, and a volumetric water content at field capacity of 39.2% (Soil Analysis Laboratory, INRA Arras, France). It contains 1.5% total organic matter, 0.84% carbon, 0.1% nitrogen, with a C:N ratio of 8.4. The mesocosm set-up aimed at reproducing the agricultural field by using its soil and adding locally present earthworms and a plant species commonly used as a model system for cereal grass.

Adult individuals of *Lumbricus terrestris* L. were supplied by Wurmwelten company (Dassel, Germany) and weighed on average 4.8 ± 1.3 g fresh weight. Adult individuals of *Aporrectodea icterica* Savigny were harvested from a pesticide-free orchard in Avignon by manual digging and weighed on average 0.4 ± 0.2 g fresh weight. The earthworms were kept in their original soil for 3 days at a temperature of 14 ± 2 °C and then placed in a mixture of the original soil and the experimental soil for one week at 8 °C in the dark before the onset of the experiment.

*Brachypodium distachyon* L. was the plant species selected for this study due to its small size and short life cycle of less than 3 months [39] and because it is frequently used in controlled environment experiments [40]. The seeds of the wild type variety (Bd 21 WT), were supplied by Observatoire du Végétal, INRAE Versailles (Paris, France). After one week germination in seedling trays in vermiculite, four seedlings were planted in each mesocosm, outside the central cylinder that was introduced as a base for flux measurements of greenhouse gases (see Fig S1). During the first 3 weeks, any dead seedlings were replaced. The experiment finished with a final destructive harvest.

### Mesocosm design and experimental treatments

The mesocosms consisted of PVC tubes of 16 cm diameter and 37 cm height (Fig S1). Each mesocosm was filled up to 3 cm from the brim with 9.2 kg of soil already containing 10% gravimetric water content and sieved to 2 mm and compacted to a bulk density of 1.21 g cm^-3^. The mesocosms were sealed at their base with a 1 mm mesh followed by a PVC lid pierced with 5 holes (1 cm in dia.) that allowed the drainage of surplus water out of the mesocosm. A transparent plastic film of 10 cm height was attached around the top perimeter of the mesocosms in order to act as a barrier preventing the earthworms to leave the mesocosms. As a previous meta-analysis indicated that earthworm effects on plant growth are more prevalent in the presence of crop residues which serve as a food resource for soil biota [6], the soil surface of all mesocosms was covered with 4 g litter mixture (2.2% N, C/N = 24) consisting of 1.3 g dry weight of *Medicago truncatula* Gaertn. shoots and 2.7 g dry weight of *Zea mays* L. leaves, the equivalent of organic residue inputs of 1060 kg C ha^-1^ and 44 kg N ha^-1^.

The mesocosm experiment presented in this study included an earthworm treatment (henceforth Ew) with four levels: a control without earthworms, an anecic earthworm species (*L. terrestris*) with two individuals weighing on average 9.6 ± 1g fresh weight (FW) per replicate, an endogeic earthworm species (*A. icterica*) with 7 ± 1.1 individuals weighing on average 2.9 ± 0.1g FW per replicate, and a mixture of both species with one *L. terrestris* individual (4.9 ± 0.9 g FW biomass) and 5 ± 1.6 *A. icterica* individuals (1.7 ± 0.5 g FW biomass) per replicate. The earthworm FW biomass is the equivalent of 480, 145 and 330 g m^-2^ for the anecic, endogeic and both earthworm treatments levels, respectively, which is 2 to 3 fold higher than in the field of origin of the soil where earthworm total biomass varied from 98 to 135 g m^-2^. Note that *L. terrestris* was more recently re-classified as species displaying traits belonging to both anecic and epigeic species [36], but in the vast majority of literature it is considered an anecic species. The earthworm treatment was factorially crossed with a plant (*B. distachyon* L.) treatment with two levels, with and without plants, with 7 replicates per treatment combination.

The experiment was carried out in a greenhouse kept at temperatures ranging between 20 and 23°C during day-time and 18-20°C during night-time with an air relative humidity of 80%. The natural light was supplemented during day-time by artificial lighting for 12 hours per day with the help of high-pressure sodium lamps. The mesocosms were divided into two blocks corresponding to their position on the north or south bench of the greenhouse, and their position within the block was randomly changed twice a week, limiting the bias of the position within the block. The experiment run for 12 weeks between March and May 2017.

### The watering protocol

The watering protocol was specifically designed to include soil moisture fluctuations (analogous to what happens in natural conditions) and to allow the earthworm burrowing to affect soil water content (SWC). At the beginning of the experiment, the mesocosms were watered with 1.7 L of reverse-osmosis water using a laboratory dispenser (two sessions of 850 ml each), a volume sufficient to observe water draining out of the mesocosms from the pierced-lid at the bottom of mesocosms.

Measurements of weight changes after 24 h were used to calculate the weight of the mesocosms at field capacity (knowing that the soil already contained 10% gravimetric water, i.e. ∼0.9 L). Changes in total mesocosm weight hence allowed for the estimation of the soil water content during the experiment and its expression in terms of % of field capacity. At field capacity, the mesocosms Water Filled Pore Space (WFPS) can be estimated at 71.0± 2.5% on average (calculated as WFPS = water content/porosity; porosity = 1-bulk density/particle density; particle density = 2.7 Mg m^-3^). The mesocosms were exposed to a drying phase until one of them reached almost 50% of field capacity, which happened after 6 weeks. The volume of water lost by the driest mesocosm was determined and then added (second watering in two sessions again) to all mesocosms to set them back to 100% of field capacity. A second drying phase was imposed until the penultimate week where a third watering was done with the same amount of water as for the second watering. Following this method, all mesocosms experienced two drying-rewetting cycles (Fig 1A,B).

**Figure 1.**
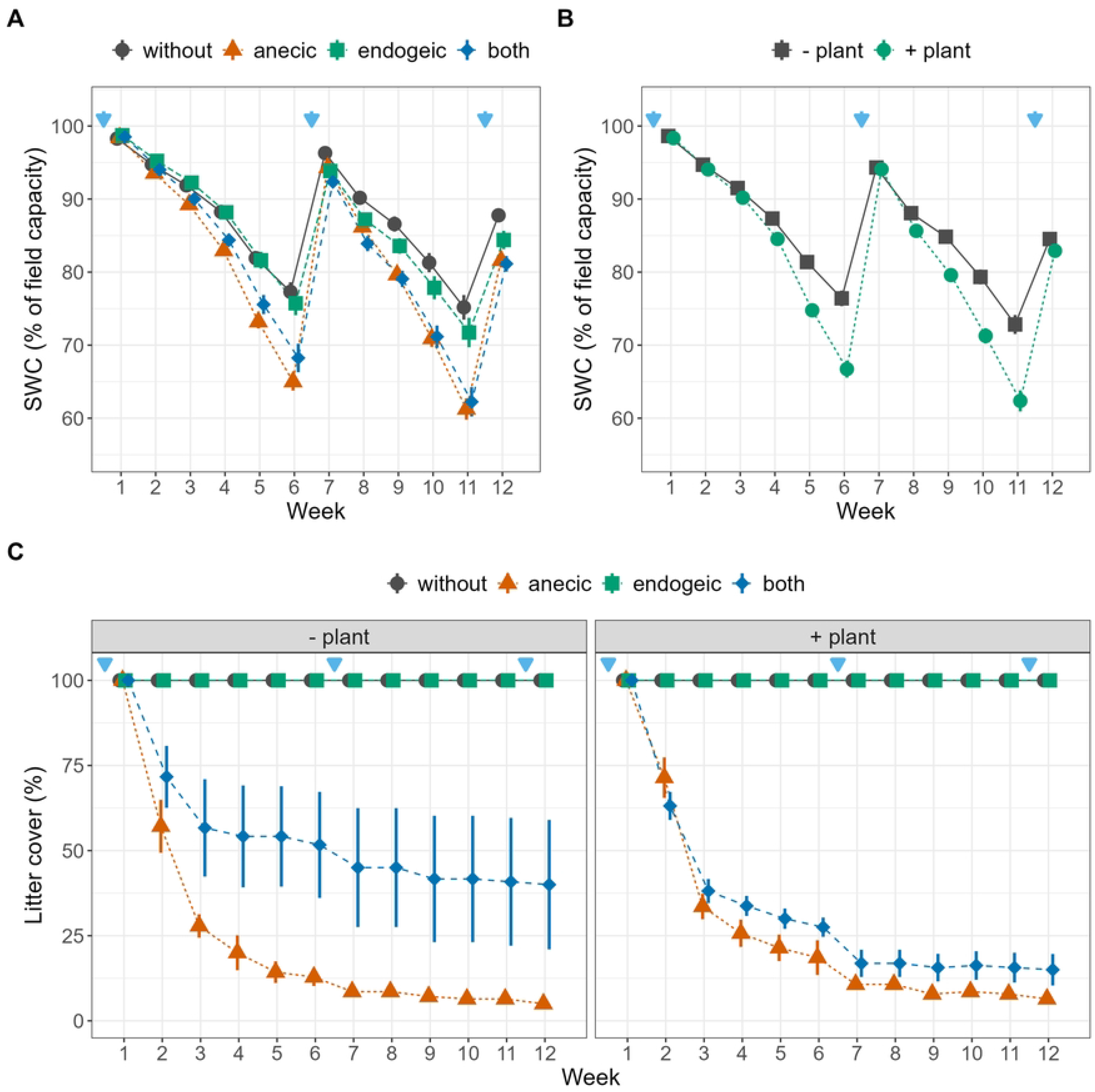
Temporal dynamic of soil water content and litter cover. (A,B) Temporal dynamic of soil water content (SWC) expressed as percentage of field capacity (C,D) and percentage of surface covered by litter as affected by the earthworm and plant treatment. Blue arrows represent watering events. Error bars represent ± 1 SEM. Different letters represent significantly different levels as estimated by Tukey’s HSD post hoc test. The effects of earthworm treatment independent of plant treatment, and vice-versa, are displayed when the Ew×Plant interaction is not significant (A,B).

### Response variables

CO_2_ and N_2_O emissions measurements were carried out weekly, and during the weeks with watering, always 24h after watering events, in order to capture eventual emission peaks. A static sampling chamber approach was used, and following the recommendations of Rochette (2011) [41]. The sampling chamber (9 cm dia., 6 cm height, 370 ± 1 mL, Fig S1) was equipped with a bung of silicone rubber for gas sampling at the top. During sampling, the chamber was placed on the circular collar (Fig S1) that was inserted in the center of the mesocosms during the setup of the mesocosms, and which allowed measuring soil N_2_O and CO_2_ fluxes without disturbing the plants. The collar was inserted into the soil down to 3 cm and consisted of frame that provided support but allowed the access by earthworms and roots to the inner soil core thanks to two opening windows. The aboveground part of the collar contained a gutter-like double walled-section/groove where the static chamber was placed during sampling. On the day of gas sampling, the mesocosms were transported to the gas sampling laboratory with a trolley, the two block being measured over two days. Prior to sampling, 20 ml of distilled water was added in the groove to provide air-tight sealing when placing the static chamber on the collar. CO_2_ and N_2_O fluxes were measured at the Platform for Chemical Analysis in Ecology (LabEx CeMEB, Montpellier, France). CO_2_ concentrations were measured with a gas chromatograph (MicroGC S-Series, SRA 9 Instruments, Marcy l’Etoile, France) using a catharometric detector, quantifying the gases on the basis of their thermal conductivity. N_2_O concentrations were measured by gas chromatography equipped with an electron capture detector (Varian CP-3800, Varian Inc., Palo Alto, USA). Air samples were taken at T0 (immediately after placing the chamber on the collar) and after 2 h to assess the changes in CO_2_ and N_2_O concentrations. Previous tests sampling after 1, 2, 3 and 4 hours revealed that gas accumulation was linear during this short time. A volume of 0.2 mL was sampled sequentially for gas measurements via the silicone bung using a plastic syringe equipped with a 25G needle and injected immediately in the gas chromatographs via a 1/32“ PFA line. Concentration changes in the sampling chamber between T0 and T0+2h were used to estimate the greenhouse gas emission rates and converted as g C-CO_2_ (or N-N_2_O) m^-2^ day^-1^.

At the end of the experiment, the mesocosms were transported to the INRAE center of Nancy to analyze soil macroporosity by X-ray tomography using a medical scanner (BrightSpeed Exel 4, General Electric), with settings of 120 kV and 50 mA for the current and 0.625 mm width for each image. Images were transformed into 16-bit image and binarized (i.e. converted in black and white) using a fixed threshold value [42] since the different peaks (for the soil matrix and for the porosity) were well separated [43]. Roots and associated pores could not be included in the analysis due to their smaller average size compared to the resolution of the scanner (0.4 mm per pixel). The burrow system was then characterized by computing the volume and number of burrows in four soils layers (L1 for 0-8.5 cm, L2 for 8.5-17 cm, L3 for 17-25.5 cm and L4 for 25.5-34 cm depth, Fig S2) using ImageJ [33]. Drying- rewetting cycles contributed to the formation of cracks, i.e. macropores resulting from physical processes (shrinkage, swelling [33]) notably in the top soil layer (Fig S2). We attempted to differentiate cracks from burrows according to the macropore circularity in 2D images, since earthworm burrows are more circular compared to cracks. However, this method still identified burrows in the control mesocosms (without earthworms), with 25.4% of pores misidentified as burrows in the whole mesocosm, and higher error in the first layer (32.2%) than in the bottom layer L4 (20.1%) (Fig S2). Therefore, assuming that gas fluxes are influenced by the total porosity, regardless of its biological or physical origin, and the high correlation between the 15 different porosity variables (Fig S3), although we investigated how the treatment affected the different type of soil pore (pores, burrows, and cracks), we decided to only use the total pore volume data (burrows and cracks) as a predictor in the models.

After the X-ray scan in Nancy the mesocosms were transported back to Montpellier for the final destructive harvest. The proportion of earthworms found at the final harvest was 62 and 90% for *L. terrestris* and *A. icterica,* respectively. The proportion of recovered earthworms was likely affected by the mortality occurring during the days of transport and storage for the X-ray scans (during which the temperatures and vibrations were not controlled) as several *L. terrestris* individuals were found freshly dead at the harvest. However, the X-ray scans together with the litter mass loss dynamic (Fig 1C) provide strong evidence that the earthworms were active during the whole duration of the experiment. Litter cover was assessed weekly by the same person using a visual estimation method with 5% intervals. Other additional soil and plant-related response variables were measured at the end of the experiment (Fig S3). Soil analyses were performed on an homogenized soil sample from the upper 10 cm of the mesocosms inside the collar and sieved at 2mm. Potential soil microbial denitrification enzymatic activity (DEA) was measured using the acetylene inhibition method described by a method that measures total potential denitrification (as N_2_O and N_2_) [44]. This is a complementary method to the fluxes measured during the experiment which are only measuring the N_2_O emissions. The MicroResp^TM^ method was used to determine the microbial metabolic quotient [45]. Approximately 0.39 g dry weight of soil was incubated in six replication wells with a solution of D-glucose (1.5 mg C g^-1^ soil), and six replication wells with deionized water (for basal respiration), so as to reach 80% of the field capacity in 96-DeepWell Microplates (Fisher Scientific E39199). Cresol red gel detection plates were prepared as recommended by the manufacturer. After an initial two-hour pre-incubation at 25°C in the dark, each deepwell microplate was covered with a CO_2_-trap microplate detection plate using a silicone gasket (MicroResp™, Aberdeen, UK). The assembly was secured with a clamp, and incubated for four additional hours. Optical density at 590 nm (OD590) was measured for each detection well before and after incubation using a Victor 1420 Multilabel Counter (Perkin Elmer, Massachusetts, USA). Calibration relying absorbance (OD590 readings) and CO_2_ concentrations was performed using the gas chromatograph previously described. Final OD590 were normalized using pre-incubation OD590, and converted as respiration rates expressed in µg C-CO_2_ respired per g^-1^ of soil per h^-1^. The glucose-induced respiration rate was used to estimate the soil microbial C (C_mic_, µg C_microbial_ g^-1^ dry soil) biomass [46]. Finally, the metabolic quotient (Met_Q) was determined as the ratio between basal respiration rates measured in the wells with water only and no C substrate (as a proxy of the microbial basal respiration) and C_mic_. Soil mineral nitrogen was extracted from 10 g of freshly sampled soil with 40 mL of 1 M KCl solution. Nitrate and ammonium concentrations were measured by continuous flow spectrophotometry (SKALAR 3000 auto analyzer, Breda, The Netherlands). The plant shoot biomass was weighed after drying at 60°C for three days. As the roots of *B. dystachyon* are extremely thin and fragile, it was not feasible to sample root biomass.

### Statistical analyses

Statistical analyses were done using R version 4.0.2 (R Development Core Team, 2015) in Rstudio version 1.3.959 (RStudio Team, 2015). Weekly time series of CO_2_ and N_2_O emissions and SWC were analyzed with the “nmle” package version 3.1-145 [47] to perform repeated measurers analyses using a generalized mixed-effects model to test the effect of earthworms, plants, SWC (for GHG emissions only) and sampling week and their interactions on gas fluxes. The identity (ID) of the mesocosm and its position in the blocks were used as random factors to account for temporal pseudoreplication and the effect of the position in the north or south bench in the greenhouse (“random = ∼ 1 | Block / ID”). To reach models which respect the assumption of homoscedasticity of the residuals, we tested the model fit with varIdent (for plant and earthworm experimental treatments), varPower (for SWC) and varExp (for SWC) weighting functions [48] and selected the most appropriate models based on maxim likelihood (ML) model comparison tests. A similar approach was used for the cumulative CO_2_ and N_2_O data at the end of the experiment (estimated assuming constant emission rates between the weekly measurements), but without the sampling week among the fixed effects and the mesocosm ID in the random effects. For the later analysis, we used the mean of weekly SWC values as arguably this variable is more relevant to the cumulative fluxes. The “r.squaredGLMM” function from the MuMIn package [49] was used to derive the proportion of the variation that was explained by the fixed factors (i.e. marginal r^2^, mr^2^) in mixed-effects models. The “multcom” package was used to perform Tukey’s HSD (honestly significant difference) multicomparison posthoc test, but note that this test does not include the random effects and occasionally the results are not entirely in line with the fitted coefficients from the mixed effects models.

Additional analyses were effectuated aiming to link the multiple potential predictors (Fig S3) measured at the end of the experiment and the CO_2_ and N_2_O fluxes from week 12 (just before the experiment was stopped). Multiple response variables were measured in order to explore potential predictors, a method of best subset selection that is penalizing model complexity (i.e. regularization) during estimation was required. Regularization aims to significantly reduce the variance of the model as well as model overfitting by varying the lambda (λ) parameter which is tuning the level of penalization for the complexity of the model. This approach has been proved to be a viable option for estimating parameters in scenarios with small sample size and many collinear/correlated predictors. Here we used a penalized regression method based on the minimax concave penalty (MCP) in order to select the best subsets [50] using the *ncvreg* package 3.11-1 [51]. This approach was combined with a 10-fold cross-validation procedure to derive the lambda parameter (also called the regularization rate) which minimizes the cross-validation error. We report the fitted coefficients and the coefficient of determination (r^2^) at lambda values that minimize the cross-validation error. The subset variables with retained non-zero coefficients were then tested in the generalized mixed-effects models which have the advantage of including random effects alongside the treatment factors.

## Results

### Soil water content

Over time, the soil water content (SWC), expressed as a percentage of field capacity, exhibited variations due to the drying-rewetting cycles and the treatments involving plants and earthworms, which were reflected in the Plant×Week and Ew×Week interactions (Table 1, Fig 1A and B). SWC was significantly lower in presence of plants during the three last weeks of both drying cycles, with 5% lower values in the absence of plants across the whole experiment (averaged over the earthworm treatments). The presence of anecic earthworms (with or without endogeic earthworms) led to significantly lower SWC relative to control during weeks three to six and eight to twelve. Averaged over the twelve weeks, SWC was 81.3% of field capacity in the presence of anecic earthworms, 81.7% in presence of both earthworm species, 85.9% with endogeic earthworms and 87.0% in the control.

**Table 1.** Time series and cumulative statistic table. Minimal adequate models for weekly time series (SWC, Litter cover, N_2_O, and CO_2_) and cumulative emissions (cN_2_O and cCO_2_) as affected by the earthworm (Ew), plant (Plant), sampling week (Week), soil water content (SWC) and their interactions. “NA” stands for non-applicable, “ns” stands for variables that were not significant (P > 0.1) and were not retained in the minimal adequate models whereas _m_r^2^ represents the marginal coefficient of determination. Figures are F-values. ***P < 0.001; **P < 0.01; *P< 0.05; ^+^P < 0.1.

| Source | SWC | Litter | N<sub>2</sub>O | cN<sub>2</sub>O | CO<sub>2</sub> | cCO<sub>2</sub> |
| --- | --- | --- | --- | --- | --- | --- |
| Ew | 46.68*** | 182.89*** | 82.97*** | 7.07*** | 2.53<sup>+</sup> | 1.25 |
| Plant | 173.83*** | 2.91<sup>+</sup> | 34.58*** | 3.82<sup>+</sup> | 5.98* | ns |
| Week | 2758.70*** | 68.85*** | 112.9*** | NA | 230.28*** | NA |
| SWC | NA | NA | 0.04 | 6.35* | 128.85*** | 7.54** |
| Ew×Plant | 2.63<sup>+</sup> | 3.65* | 9.52*** | 2.05 | 2.01 | ns |
| Ew×Week | 9.40*** | 198.63*** | 5.61*** | NA | 1.97** | NA |
| Plant×Week | 24.01*** | ns | 9.32*** | NA | 4.93*** | NA |
| Ew×SWC | NA | NA | 14.84*** | 4.45** | 5.16** | 3.16* |
| Plant×SWC | NA | NA | 1.43 | ns | ns | ns |
| Week×SWC | NA | NA | 8.34*** | NA | ns | NA |
| Ew×Plant×Week | ns | ns | 1.57* | NA | ns | NA |
| Ew×Plant×SWC | NA | NA | 9.51*** | ns | ns | ns |
| Ew×Week×SWC | NA | NA | 3.81*** | NA | ns | NA |
| Plant×Week×SWC | NA | NA | 5.52*** | NA | ns | NA |
| Ew×Plant×Week×SWC | NA | NA | ns | NA | ns | NA |
| $m^2$ | 0.81 | 0.91 | 0.67 | 0.49 | 0.43 | 0.27 |

### N_2_O and CO_2_ weekly and cumulative fluxes

Weekly N_2_O emissions were significantly affected by all possible three-way interactions between earthworms, plants, SWC and time (Table 1, Fig 2A). N_2_O emissions were higher after each watering (with the measurements always being done 24h after watering), and were the highest in the second week of the experiment (0.10 g N-N_2_O m^-2^ day^-1^, Fig 2A). The intensity and duration of these emission peaks depended on the earthworm and plant treatments, but varied with time, as indicated by the significant Ew×Plant×Week interaction. For example, N_2_O emissions peaked ten days after the first watering in presence of anecic earthworms compared to in presence of endogeic earthworms, and reached the maximum in the absence of plants. Cumulative N_2_O emissions over the 12 weeks of the experiment were the highest in the control without earthworms or plants (0.135 g m^-2^, Fig 2B,C).

**Figure 2.**
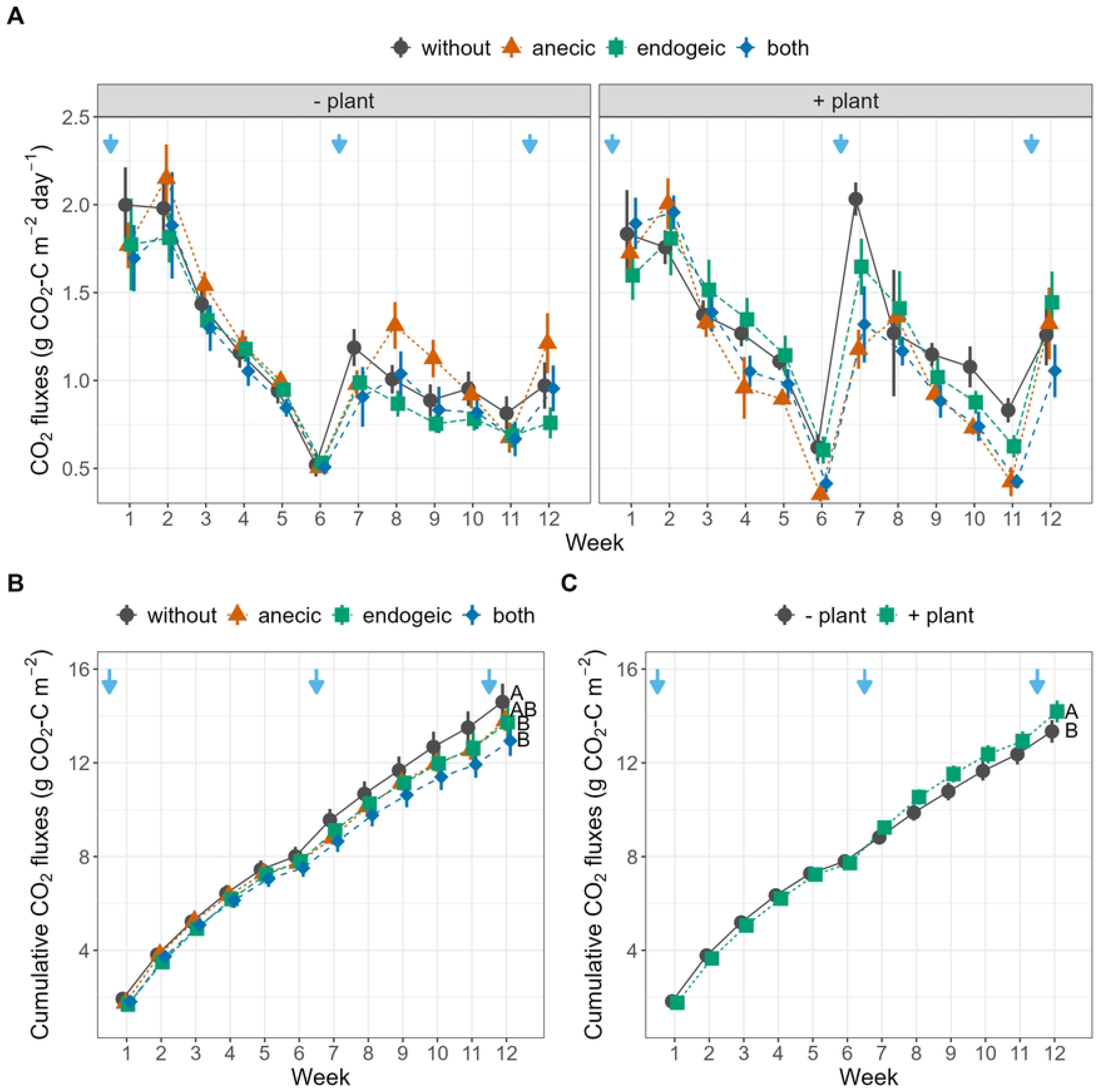
N_2_O emissions through the 12 weeks experiment. (A,B) Weekly (C,D) and cumulative N-N_2_O emissions as affected by the earthworm and plant treatment. Blue arrows represent watering events. Error bars represent ± 1 SEM. Different letters represent significantly different levels as estimated by Tukey’s HSD post hoc test. The effects of earthworm treatment independent of plant treatment, and vice-versa, are displayed when the Ew×Plant interaction is not significant (B, C).

Relative to control, the cumulative N_2_O emissions were 17.0, 34.6 and 44.8% lower in the anecic, both and endogeic treatments, respectively, and 19.8% lower in presence of plants (Fig 2C,D). The Ew×SWC interaction indicates that cumulative N_2_O emissions increased with average SWC for control, anecic and both earthworm treatments, but decreased with SWC in the presence of endogeic earthworms (Table 1, Fig 4A). CO_2_ weekly emissions were significantly affected by the Ew×Week, Plant×Week and Ew×SWC interactions and were higher after re-watering at higher SWC (Table 1, Fig 3A). The cumulative CO_2_ emissions showed no significant response to earthworm or plant presence (Fig 3B,C). However, a significant interactive effect of earthworms with SWC was found (Table 1, Fig 4E) indicating that when SWC was relatively high (> 85.9% SWC on average), as in the endogeic and earthworm-control treatment, cumulative CO_2_ emissions generally decreased with increasing SWC. The opposite was true in the presence of anecic, while no relationship was found in the presence of both species. Additionally, when the SWC effect is not included as a predictor, the cumulative CO_2_ emissions relative to control were 5.9% and 11.4% lower in the endogeic and both earthworm treatment respectively as indicated by the Tukey HSD post hoc test (Fig 3B). Conversely, plant presence increased the cumulative CO_2_ emissions by 6%.

**Figure 3.**
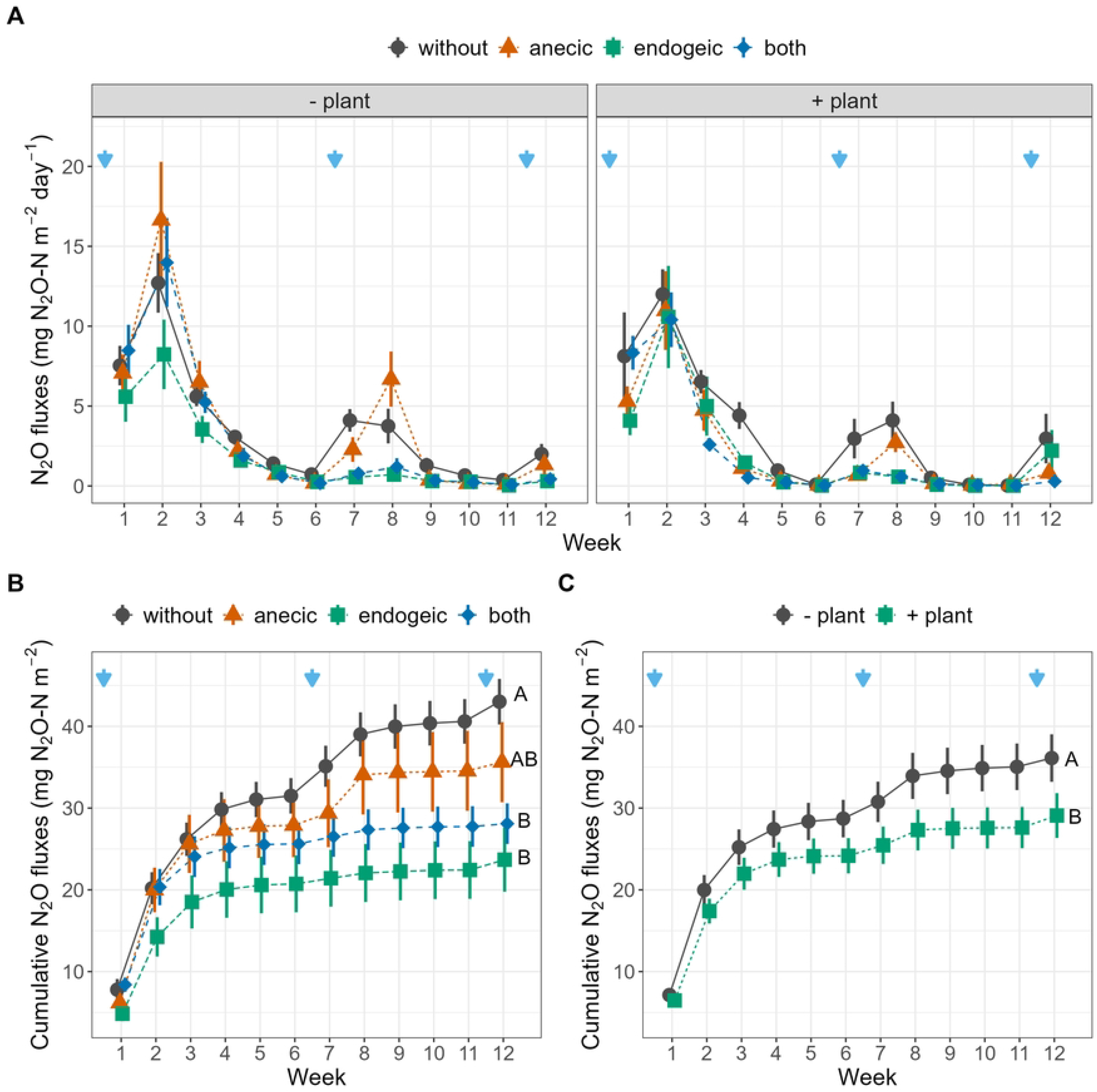
CO_2_ emissions through the 12 weeks experiment. (A,B) Weekly (C,D) and cumulative C-CO_2_ emissions as affected by the earthworm and plant treatment. Blue arrows represent watering events. Error bars represent ± 1 SEM. Different letters represent significantly different levels as estimated by Tukey’s HSD post hoc test. The effects of earthworm treatment independent of plant treatment, and vice-versa, are displayed when the Ew×Plant interaction is not significant.(B, C).

**Figure 4.**
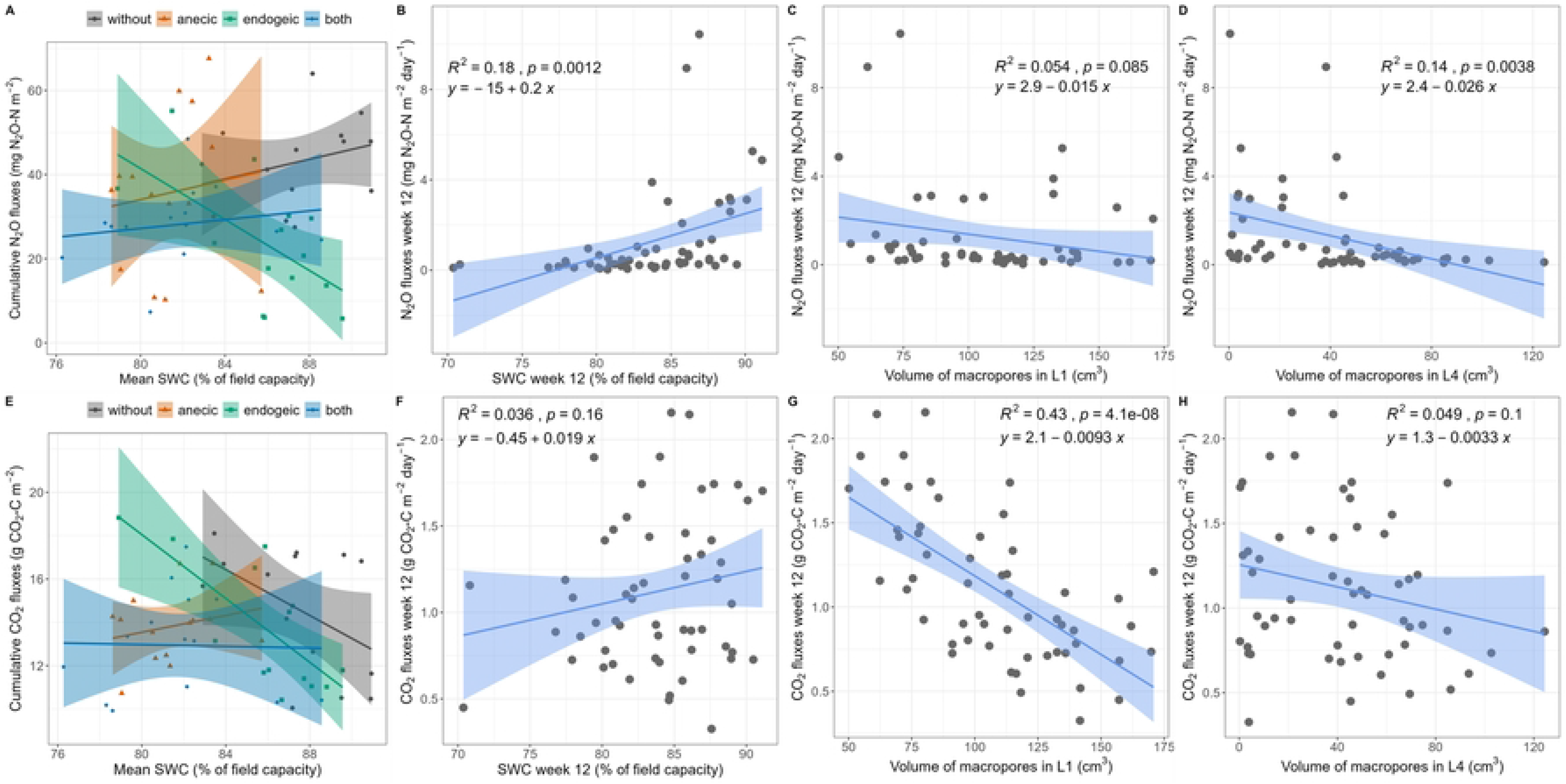
N_2_O and CO_2_ emissions as affected by soil water content and macroporosity. (A,E) total cumulative emissions as affected by the Ew×SWC interaction, where SWC represents the 3- month average SWC. (B, C, D, F, G, H) Relationships between N_2_O and CO_2_ emissions measured in the last experimental week (week 12), (C, F) soil water content and (D,G) X-ray tomography estimated volume of macropores in the top-soil layer (L1), and (D, H) in the bottom layer for N_2_O (L4), When significant, linear regression line and 95% confidence intervals are displayed along with regression equation, coefficient of determination (*R²*) and p-value. Note that this relationship may differ from the mixed-effect models results (Table 1 and 4).

### Plant, litter, microbial activity, nutrient and porosity

Although not significant, the presence of earthworms led to higher aboveground plant biomass compared to control (+57, +25 and +41% for anecic, endogeic and both species, respectively, Table 2, Fig S4A). Litter cover depended on the presence of earthworms and varied with time and plant presence as indicated by the Ew×Week and Ew×Plant interactions respectively (Table 1, Fig 1C). After

**Table 2.** Covariables summary statistics table. Summary statistics (mean ± standard error) and minimal adequate models for the covariables considered as predictors for the N_2_O and CO_2_ emissions from the last week of the experiment in addition to the experimental treatments. Different letters represent significantly different treatments according to Tukey’s HSD post hoc test. F-values are shown with significance levels: ***P < 0.001; **P < 0.01; *P< 0.05; +P < 0.1. _m_r^2^ = marginal coefficient of determination . Ew= “earthworm”, DEA= denitrifying enzyme activity, BR= basal respiration, Met_Q) metabolic quotient.

| Plant | Earthworm | Total shoot biomass<br>dry g | Litter cover<br>% | DEA<br>g N g soil<sup>-1</sup> h<sup>-1</sup> | Cmic<br>g C-CO<sub>2</sub><br>g soil<sup>-1</sup> h<sup>-1</sup> | BR<br>g C-CO<sub>2</sub><br>g soil<sup>-1</sup> h<sup>-1</sup> | Met_Q<br>Ratio | NO<sub>3</sub><sup>-</sup><br>mg kg<sup>-1</sup> | NH<sub>4</sub><sup>+</sup><br>mg kg<sup>-1</sup> |
| --- | --- | --- | --- | --- | --- | --- | --- | --- | --- |
| - plant | without |  | 100 ± 5.6 c | 0.04 ± 0.01 ab | 56.9 ± 2.5 a | 0.8 ± 0 a | 0.5 ± 0 a | 31.7 ± 5.9 abc | 0.4 ± 0.1 a |
| + plant | without | 3.6 ± 0.8 a | 100 ± 6.5 c | 0.05 ± 0.01 ab | 62.3 ± 2.9 a | 0.8 ± 0.1 a | 0.5 ± 0 a | 12.9 ± 6.8 a | 0.3 ± 0.1 a |
| - plant | anecic |  | 5 ± 6 a | 0.07 ± 0.01 ab | 63 ± 2.6 a | 0.8 ± 0.1 a | 0.5 ± 0 a | 54.6 ± 6.3 cd | 0.4 ± 0.1 a |
| + plant | anecic | 5.6 ± 0.8 a | 6.4 ± 6 a | 0.05 ± 0.01 ab | 58.9 ± 2.6 a | 0.9 ± 0.1 a | 0.5 ± 0 a | 32.4 ± 6.3 abc | 0.2 ± 0.1 a |
| - plant | endogeic |  | 100 ± 6 c | 0.04 ± 0.01 a | 54.4 ± 2.6 a | 0.8 ± 0.1 a | 0.6 ± 0 a | 49.1 ± 6.3 bcd | 0.2 ± 0.1 a |
| + plant | endogeic | 4.5 ± 0.8 a | 100 ± 6 c | 0.05 ± 0.01 ab | 59 ± 2.6 a | 0.8 ± 0.1 a | 0.5 ± 0 a | 23.1 ± 6.3 ab | 0.2 ± 0.1 a |
| - plant | both |  | 40 ± 6.5 b | 0.06 ± 0.01 ab | 59.7 ± 2.9 a | 0.8 ± 0.1 a | 0.5 ± 0 a | 65.9 ± 6.8 d | 0.2 ± 0.1 a |
| + plant | both | 5 ± 0.7 a | 15 ± 5.6 ab | 0.07 ± 0.01 b | 61.3 ± 2.5 a | 0.9 ± 0 a | 0.5 ± 0 a | 39.2 ± 5.9 abcd | 0.4 ± 0.1 a |
| Source |  | Statistical results (minimal adequate models) |  |  |  |  |  |  |  |
|  | Ew | ns | 74.39*** | 2.75<sup>+</sup> | ns | ns | ns | 15.95*** | 2.22<sup>+</sup> |
|  | Plant | NA | 0.93 | ns | ns | ns | ns | 46.41*** | ns |
|  | Ew×Plant | NA | 3.08* | ns | ns | ns | ns | ns | ns |
|  | m<sup>2</sup> | 0.0 | 0.91 | 0.14 | 0.00 | 0.00 | 0.00 | 0.73 | 0.11 |

12 weeks, anecic earthworms alone reduced litter cover to 5 And 6.4% in absence and presence of plant respectively and, when both species were present, to 40 and 15% in absence and presence of plant respectively (Fig S4B). At week 12, SWC was positively correlated with the litter cover (r = 0.55, t = 4.86, P < 0.001, n = 56; Fig S3), suggesting that, at least in part, the presence of earthworms (especially the anecic *L. terrestris*) also affected SWC via higher evaporation from bare soil due to litter burial. We found marginally higher denitrification potential in presence of anecic earthworms (+22%) relative to control, but no plant effects (Table 2, Fig S4C). Soil nitrate content (NO_3_ ^-^) was always lower (-80%) in the presence of plants and increased in the presence of earthworms with synergistic effects in presence of both species (Table 2, Fig S4G). Macropore volume in top soil layer (L1) was not affected by any experimental treatment (Table 3, Fig S3). We found lower macropore volume in the presence of plants in the other three layers (-26.0% in L2, -23.8% in L3, -18.1% in L4). Macropore volume in L2-L4 were affected by the earthworm treatment, with the highest volume in the mesocosms with endogeic earthworms, followed by both, anecic and control treatment (Fig S3). Similar trends were observed when macropores were differentiated into burrows and cracks (Table S1, Fig S3). Burrow volume was largely driven by the earthworm treatment with high coefficients of determination in L2, L3, and L4 and in total (i.e. the sum of L1 to L4). Plant presence only affected total burrow volume and in L2 (Table S1).

**Table 3.** Covariables summary statistics table. Summary statistics (mean ± standard error) and minimal adequate models of the porosity variables considered as predictors for the N2O and CO2 emissions from the last week of the experiment in addition to the experimental treatments. Different letters represent significantly different treatments according to Tukey’s HSD post hoc test. F-values are shown with significance levels: ***P < 0.001; **P < 0.01; *P< 0.05; +P < 0.1. _m_r^2^ = marginal coefficient of determination. Ew= “earthworm”,

| Plant | Earthworm | Vpores_L1<br>cm<sup>3</sup> | Vpores_L2<br>cm<sup>3</sup> | Vpores_L3<br>cm<sup>3</sup> | Vpores_L4<br>cm<sup>3</sup> | Vpores_tot<br>cm<sup>3</sup> |
| --- | --- | --- | --- | --- | --- | --- |
| - plant | without | 127.1 ± 10.9 a | 79.7 ± 5.9 bc | 39.4 ± 5.7 ab | 9 ± 5.6 a | 255.2 ± 23 abc |
| + plant | without | 98.8 ± 12.6 a | 57.4 ± 6.8 ab | 20.3 ± 6.6 a | 2.1 ± 6.5 a | 178.6 ± 26.6 a |
| - plant | anecic | 96.9 ± 11.7 a | 60.3 ± 6.3 ab | 57.6 ± 6.1 bc | 39.3 ± 6 bc | 254.1 ± 24.6 abc |
| + plant | anecic | 99.3 ± 11.7 a | 45.5 ± 6.3 a | 37.7 ± 6.1 ab | 23.5 ± 6 ab | 206 ± 24.6 ab |
| - plant | endogeic | 118.4 ± 11.7 a | 96.6 ± 6.3 c | 92 ± 6.1 d | 77.3 ± 6 d | 384.2 ± 24.6 d |
| + plant | endogeic | 92.6 ± 11.7 a | 71.8 ± 6.3 abc | 72.2 ± 6.1 cd | 63.5 ± 6 cd | 300 ± 24.6 bcd |
| - plant | both | 105.6 ± 12.6 a | 77.3 ± 6.8 bc | 78.8 ± 6.6 cd | 65.7 ± 6.5 cd | 327.4 ± 26.6 cd |
| + plant | both | 112.5 ± 10.9 a | 55.1 ± 5.9 ab | 63.2 ± 5.7 bc | 53.6 ± 5.6 cd | 284.4 ± 23 abcd |
| Source |  | Statistical results (minimal adequate models) |  |  |  |  |
| Ew |  | ns | 9.04*** | 32.04*** | 191.71*** | 11.91*** |
| Plant |  | ns | 26.63*** | 21.36*** | 13.83*** | 15.78*** |
| Ew×Plant |  | ns | ns | ns | ns | ns |
| m<sup>2</sup> |  | 0.00 | 0.48 | 0.68 | 0.94 | 0.59 |

### Exploration of multiple predictors for final N_2_O and CO_2_ fluxes

Out of the 16 tested potential predictors (Fig S3), the MCP-penalized multiple regression for N_2_O emissions indicate that, in addition to a retained positive coefficient for SWC (4.663e^-04^), the macropore volume in the first and fourth soil layer (Vpores_L1 and Vpores_L4) were also retained with negative coefficient (-1.01e^-05^ and -4.18e^-05^ respectively) at minimum cross validation error with lambda = 0.001 (Fig 4 A to D). CO_2_ emissions were influenced by macropore volume in the top soil (Vpores_L1), with a negative coefficient (- 0.034) at minimum cross validation error with lambda = 0.321 (Fig 4 E to H). These selected predictors also had among the highest correlation coefficients with CO_2_ and N_2_O fluxes (Fig S2) as shown by univariate regression. Finally, we compared how the inclusion of porosity metrics affected models’ performance in explaining the emissions from the last experimental week (Table 3). For N_2_O emissions, Vpores_L1 and Vpores_L4 could not be included together due to overfitting and convergence issues, leading us to ran separate model for each porosity variable.

Minimal adequate models for N_2_O emissions including Vpores_L1 explained more variation than without the porosity metric (r²m = 0.86, vs. r²m = 0.69, respectively) and retained two additional interactions, notably Ew×SWC×Vpores_L1 three-way interaction with a positive fitted coefficient (Table 4). The second model including Vpores_L4 had an intermediate amount of explained variation (r^2^m = 0.72), and only detected one additional marginally significant SWC×Vpores_L4 interaction (p-value = 0.054, Table 4). Regarding CO_2_ emissions, including Vpores_L1 largely increased the amount of variance explained (r²m = 0.83 vs. r²m = 0.18, Table 4). The four-way interaction Ew×Plant×SWC×Vpores_L1 was significant with a positive fitted coefficient, indicating higher CO_2_ emissions in the presence of earthworms (all treatment combinations containing earthworms) and plants, under high SWC levels and high macropore volume in L1.

**Table 4.**
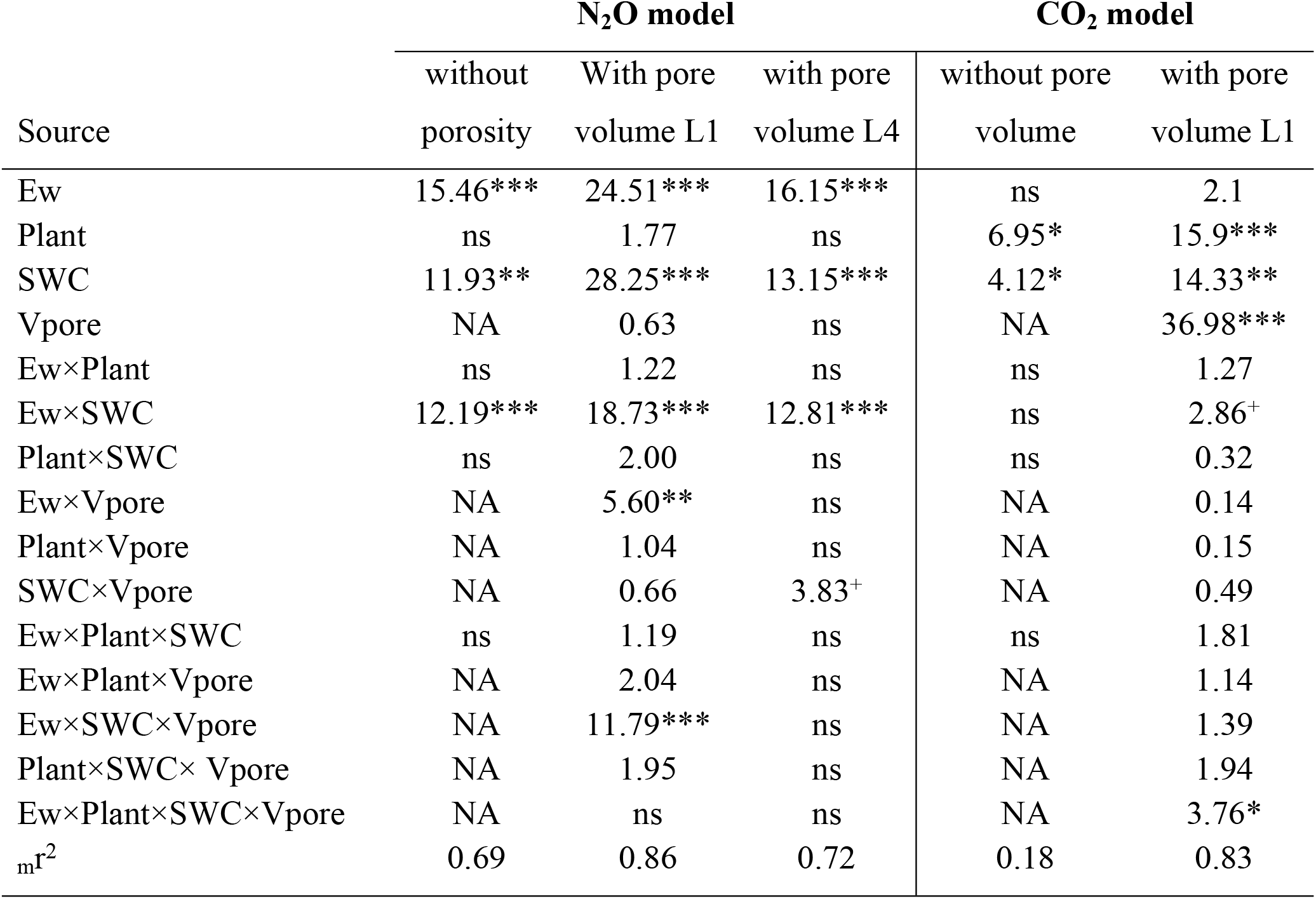
Minimal adequate models presenting the results explaining the CO_2_ and N_2_O fluxes from the last sampling (week 12) where the soil porosity-related variables were included in the model as potential predictors (compared with the models without the soil-porosity variables). “NA” stands for non-applicable, “ns” stands for variables that were not significant and were not retained in the minimal adequate models and _m_r^2^ represents the marginal coefficient of determination. F-values are shown with significance levels: ***P < 0.001; **P < 0.01; *P< 0.05; ^+^P < 0.1.

| Source | N<sub>2</sub>O model |  |  | CO<sub>2</sub> model |  |
| --- | --- | --- | --- | --- | --- |
|  | without porosity | With pore volume L1 | with pore volume L4 | without pore volume | with pore volume L1 |
| Ew | 15.46*** | 24.51*** | 16.15*** | ns | 2.1 |
| Plant | ns | 1.77 | ns | 6.95* | 15.9*** |
| SWC | 11.93** | 28.25*** | 13.15*** | 4.12* | 14.33** |
| Vpore | NA | 0.63 | ns | NA | 36.98*** |
| Ew×Plant | ns | 1.22 | ns | ns | 1.27 |
| Ew×SWC | 12.19*** | 18.73*** | 12.81*** | ns | 2.86<sup>+</sup> |
| Plant×SWC | ns | 2.00 | ns | ns | 0.32 |
| Ew×Vpore | NA | 5.60** | ns | NA | 0.14 |
| Plant×Vpore | NA | 1.04 | ns | NA | 0.15 |
| SWC×Vpore | NA | 0.66 | 3.83<sup>+</sup> | NA | 0.49 |
| Ew×Plant×SWC | ns | 1.19 | ns | ns | 1.81 |
| Ew×Plant×Vpore | NA | 2.04 | ns | NA | 1.14 |
| Ew×SWC×Vpore | NA | 11.79*** | ns | NA | 1.39 |
| Plant×SWC× Vpore | NA | 1.95 | ns | NA | 1.94 |
| Ew×Plant×SWC×Vpore | NA | ns | ns | NA | 3.76* |
| $m^2$ | 0.69 | 0.86 | 0.72 | 0.18 | 0.83 |

## Discussion

To further advance our understanding of the effects of earthworms on GHG emissions, our study was designed to simultaneously investigate the effect of earthworms, plants, soil moisture fluctuations, and their interactions, with an experimental set-up allowing earthworms and plants to affect soil water status and macroporosity. In line with our first hypothesis, we found not only that the presence of earthworms did not increase the CO_2_ and N_2_O cumulative emissions over 12 weeks, but also that the presence of the endogeic species *A. icterica* (alone, or with the anecic *L. terrestris*), and the presence of plants reduced N_2_O cumulative emissions. Furthermore, earthworms, plants, and their interaction modulated soil water content fluctuations and jointly affected weekly N_2_O and CO_2_ emissions. We finally found that GHG emissions were partly explained by increased macropore volume in the first soil layer, due to earthworm burrowing activity.

By imposing soil moisture fluctuations and allowing earthworms and plants to modulate these fluctuations, our results illustrate how soil water availability controls N_2_O and CO_2_ emissions in complex ways [52,53]. In general, a combination of limited substrate diffusion at very low water content and limited gas diffusion at high water content lead to maximal N_2_O emissions (via nitrification and denitrification) and CO_2_ emissions (via respiration) at intermediate soil water content, around 75% of water-filled pore space [18,19,54]. In our experiment, SWC varied considerably, being lowest in the presence of plants and anecic earthworms, alone or mixed with endogeic earthworms (Fig 1A,B). The significant Ew×SWC interaction (Table 1) observed for cumulative and weekly N_2_O and CO_2_ emissions illustrate the SWC optimal value phenomenon which occurred around 85% of field capacity, or 59% of water filled pore space (Fig 4A,B). Indeed, anecic earthworms create large vertical burrows increasing water infiltration, and burry leaf litter increasing the proportion of bare soil and water evaporation [32]. This led to lower than optimum SWC value for microbial activity, hence the positive relationship between SWC and respiration for this species. Conversely, the treatment combinations with the endogeic species, similar to the control with no earthworms, maintained on average higher than optimal SWC for soil respiration, thus explaining the negative slopes of CO_2_ fluxes with increasing SWC. Finally, the presence of both earthworm species led to SWC values spanned across the optimum and no clear relationship between SWC and CO_2_ could be detected. Simultaneously, the presence of *B. dystachion* grass lowered the average SWC compared to the mesocosms without plants (mean ± se = 82.0 ± 0.6% of WHC) and in this soil moisture range CO_2_ emissions increased with SWC (Fig S5). In the absence of *B. dystachion,* the average SWC was higher (86.2 ± 0.6% of WHC), but increasing SWC lowered CO_2_ emissions as the range of SWC was beyond optimum, and presumably soil respiration was limited by O_2_ diffusivity under these conditions (Fig S5). Regarding N_2_O emissions, we observed similar patterns in most cases, except for the control group without earthworms, where emissions still exceeded the optimal SWC value observed in the treatments with earthworms. This finding only partially supports the hypothesis of an optimal SWC mechanism and its effect on N2O emissions. It suggests that while the interactions between earthworms, plants, and SWC strongly influence N2O emissions, other factors likely come into play when earthworms are absent. One such factor could be a significantly different soil porosity status.

The inclusion of the soil porosity data showed that the total volume occupied by soil macropores in the upper soil layer was an important predictor of GHG emissions (with a negative coefficient, Fig 4G). This suggests that increasing porosity/aeration in the top soil layer (0-8.5 cm) can decrease N_2_O and CO_2_ emissions presumably by reducing the SWC in the upper and most microbially-active soil layer. Interestingly, in line with our third hypotheses, porosity in the bottom layer (25.5-34 cm) was a good predictor of N_2_O emissions (Fig 4D) and was the only variable that is influenced by earthworm species in the same way as cumulative N_2_O emissions. Indeed, the number of burrow in the deepest layer was higher in presence of the endogeic *A. icterica* (alone or alongside the anecic species), a species with high affinity for the deepest soil layers [55], and presumably prevented the development of denitrification-stimulating anaerobic sites. The reduction of N_2_O emissions via an increased soil aeration was also previously suggested by several studies [56,57], but was not explicitly shown to our knowledge. Our results indicate that this effect is more prevalent in the presence of the endogeic species and seems to be related to the higher number of burrows that are produced by this species in contrast to the larger but less numerous semi-permanent burrows produced by the anecic species (Fig S3) [32]. In contrast, plants significantly reduced soil macropore volume, likely due to improved soil aggregate stability in the rhizosphere via roots growth and exudates [23].

Our experiment also allows to discuss the importance of nutrient availability for GHG emissions. The observed reduction of N_2_O emissions by 19.8% in presence of plants occurred likely also in part due to plant N uptake as we found that the amount of soil NO_3_ ^-^ and NH ^+^ was 43% and 20% lower respectively in the presence of plants (in line with our second hypothesis) independently of earthworm presence. This supports our hypothesis that plants can compete with microorganisms for nutrients and therefore limit the bulk microbial activity [25], given the importance of nitrogen availability for nitrification and denitrification [58]. The absence of positive effect of plants on CO_2_ fluxes (either weekly fluxes, or microbial potential activity at final harvest) is surprising, notably since our experimental design only allowed the combined measurements of CO_2_ originating from heterotrophic and root respiration. This could be explained by the nutrient (nitrate and ammonium) limitations, or by an overall low plant effect due to the relatively low plant biomass production of *B. dystachion* in our experiment. Soil NO_3_ ^-^ concentration increased in the presence of anecic and endogeic earthworms, and even more so when both ecological categories were present with, leading to a 2.1- and to a 3-fold increase relative to the control in mesocosms with and without plants, respectively. This can be explained by the combined effect of local vertical litter burial by *L. terrestris* and the horizontal redistribution in the extensive burrow system of *A. icterica* as well as higher nitrogen concentration in earthworms casts compared to bulk soil [29,59]. The accelerated burying of surface litter by the anecic species therefore likely contributed to the higher N_2_O emissions via increased N availability after watering events, but this effect faded with time and soil drying. Simultaneously, this higher nutrient availability likely contributed to higher plant growth in the presence of earthworms [6]. Despite this increase in nutrient availability, the decreased soil moisture due to earthworm burrowing and the formation of cracks in the top soil still reduced microbial activity and GHG production. Finally, the results suggest that short-term experiments are likely to emphasize this transient stimulation of emissions due earthworm bioturbation effects on organic matter mineralization, especially in experimental systems lacking plants [60–62], but also that GHG sampling mostly after watering events only, which is a common practice, will likely lead to biased estimates.

Mesocosm experiments are highly valuable tools for global change research, but care must be taken in interpreting and extrapolating results, and potential caveats of our study must be mentioned [63,64]. Since soil properties can strongly influence GHG emissions as well as earthworm casts properties [65,66], we cannot be sure of the transferability of our results to other soil types. Secondly, as gas diffusion will occur at the soil-air interface, with gases more concentrated in the soil moving towards the atmospheric air with lower concentration, the bottom of mesocosms should be as air-tight as possible, while still allowing for drainage. In our case, the holes at the bottom represented 0.4% of the surface, and were obstructed by the table, thus presumably limiting this bias. Future studies could address this issue by placing the mesocosms on a layer of the same experimental soil, or use active drainage systems consisting of suction pumps and tubing equipped with valves that can be closed after drainage. We also acknowledge that the size of our mesocosms, although larger than many other studies, could have interfered with the earthworm burrowing behavior, especially for the deep burrowing anecic earthworms [42]. Furthermore, as only one earthworm species per ecological category was used, it is unknown whether our findings are transferable to other species from the same ecological categories, or whether these findings are also valid for epigeic earthworm species which have been reported to also increase N_2_O emissions [7]. It must be noted that earthworm ecological categories were not conceptualized to describe functional role but rather ecological and morphological groups, which can also explain the high variability of earthworm species effects among the same ecological category [67]. To the best of our knowledge, our study is the first to investigate the link between earthworm-induced macroporosity and greenhouse gases fluxes, however the size of the mesocosm combined with the imposed drying-rewetting cycles led to the formation of cracks that unfortunately made the analysis more difficult. As cracks are even more unstable than burrows under drying-rewetting cycles [33], they should be avoided if possible or taken into account in analyses in future experiments. Another caveat is that our weekly measurement frequency of CO_2_ and N_2_O fluxes over 12 weeks may have missed daily variations or higher emission peaks following the watering events. However, our measurements still detected peaks after watering (Fig 2), and while stimulation of emissions under the anecic treatment was detectable, for the endogeic species, the N_2_O emission rates were consistently lower than the control. Further studies should measure emissions as frequently as possible, especially after watering events.

In conclusion, our study highlights new mechanisms by which earthworms and plants influence soil GHG emissions in an experimental set-up integrating earthworm engineering effect on soil water fluxes and soil porosity, two major mechanisms so far neglected. The presence of earthworms did not increase CO_2_ and N_2_O emissions and revealed that the endogeic earthworm *A. icterica*, a common species present in Europe and North America, even has the potential to reduce N_2_O emissions. Our study is an additional step towards a better understanding of the interaction between soil biota and soil physico-chemical properties underlying GHG emissions. Future research on these mechanisms would be highly valuable, especially in agricultural context as agriculture is the first sector of N_2_O emissions [68], and many mitigation practices are proposed (e.g. reduced tillage or cover crops) [69]. At the same time, these practices also affect earthworms communities, soil porosity, soil compaction, and water infiltration, which interact and affect soil functions [70–73]. Future research should address these points in experimental setups where the earthworm engineering effect on soil water status and aeration is allowed to take place in a realistic way.

## Acknowledgements

This study benefited from the CNRS human and technical resources allocated to the ECOTRONS Research Infrastructure. We thank Thierry Morvan for helping the organization of the soil excavation at the EFELE experimental site as well as Thierry Mathieu and David Degueldre from the technical platform Terrain d’Expériences du C.E.F.E. for their support with the greenhouse and production of gas sampling collars. Microbial and soil analyses were done at Plateforme d’Analyses Chimiques en Ecologie (PACE) supported by the LabEx CeMEB, an ANR “Investissements d’avenir” programme (ANR-10-LABX-04-01). The authors declare no conflict of interest.

## Data Availability

The datasets and R code of the current study is currently available on the first author GitHub repository at the following link https://github.com/PGanault/EarthwormPlantGHG.

## Author contributions

AM and JN provided the funding, AM, JN and PG designed the experiment, PG carried out the experiment and measurements, YC, IB, BB, AS, NF provided methods and carried out specific measurements. PG and AM analyzed the data, PG and AM wrote the paper with input from all co- authors.

